# The circadian clock gene *bmal1* is necessary for co-ordinated circatidal rhythms in the marine isopod *Eurydice pulchra* (Leach)

**DOI:** 10.1101/2022.12.21.521378

**Authors:** Lin Zhang, Edward W. Green, Simon G. Webster, Michael H. Hastings, David C. Wilcockson, Charalambos P. Kyriacou

## Abstract

Circadian clocks in terrestrial animals are encoded by molecular feedback loops involving the negative regulators PERIOD, TIMELESS or CRYPTOCHROME2 and positive transcription factors CLOCK and BMAL1/CYCLE. The molecular basis of circatidal (∼12.4 hour) or other lunar-mediated cycles (∼15 day, ∼29 day), widely expressed in coastal organisms, is unknown. Disrupting circadian clockworks does not appear to affect lunar-based rhythms suggesting a molecular independence of the two cycles. Nevertheless, pharmacological inhibition of casein kinase 1 (CK1) that targets PERIOD stability in mammals and flies, affects both circadian and circatidal phenotypes in *Eurydice pulchra (Ep)*, the speckled sea-louse. Here we show that these drug inhibitors of CK1 also affect the phosphorylation of EpCLK and EpBMAL1 and disrupt EpCLK-BMAL1-mediated transcription in Drosophila S2 cells, revealing a potential link between the positive circadian regulators and circatidal behaviour. We therefore performed dsRNAi knockdown of Ep*bmal1* as well as the major negative regulator in *Eurydice*, Ep*cry2*. Ep*cry2* and Ep*bmal1* knockdown disrupted *Eurydice*’s circadian phenotypes as expected but in addition, circatidal behaviour was also sensitive to Ep*bmal1* knockdown. Thus three *Eurydice* negative circadian regulators, EpCRY2, in addition to EpPER and EpTIM, do not appear to be required for the expression of circatidal behaviour, in contrast to the positive regulator *Ep*BMAL1. We suggest a neurogenetic model whereby the positive circadian regulators BMAL1-CLK are shared between circadian and circatidal mechanisms in *Eurydice* but circatidal rhythms require a novel, as yet unknown negative regulator.

## Introduction

Circadian clocks are composed of a number of intersected negative feedback loops in which cycling components cycle with ∼24 h rhythmicities (1,2). In higher eukaryotes such as insects and mammals these components are expressed in neurons to mediate circadian behaviour and in peripheral tissues where they control rhythmic tissue and cell-specific functions and metabolism (3,4). The core negative regulators are PERIOD, TIMELESS or CRYPTOCHROME2, which rhythmically and negatively feed back to suppress the actions of their positive transcription factors, BMAL1 (CYCLE) and CLOCK. There are also a number of kinases and phosphatases that modulate the stability of these regulators, including casein kinase 1 (CK1). *Drosophila melanogaster* has a single gene that encodes CK1, DOUBLETIME (DBT), whereas mammals have two circadian-relevant isoforms, CK1ε and CK1δ. Nevertheless, in both flies and mammals CK1 targets the stability of PERIOD proteins (5-7) and determines circadian period length (8-10), thereby highlighting a conserved function.

In contrast to circadian rhythms, the molecular bases of lunar-mediated behavioural and physiological cycles are unknown. Organisms that inhabit the intertidal zone are exposed to the gravitational pull of the moon and the sun on the oceans, so that on most coasts, high/low tide is encountered every 12.4 hours (11). Animals such as crustacea are entrained to these environmental cycles but in the absence of such tidal stimuli in the laboratory, under constant ‘free-running’ conditions, circatidal (∼12.4 h) or circalunidian (∼24.8 h) rhythms of behaviour or physiology will persist (12,13). In addition, semi-circalunar (∼15 day) and circalunar (∼29 day) rhythms have been observed in a number of intertidal organisms in which life cycle events such as spawning, emergence or reproduction are studied (14,15). Interactions between the circatidal and circadian clock have also been observed, for example, in determining the period length of marine isopod semi-lunar foraging rhythms (16).

Two main competing hypotheses that attempt to explain how circatidal behaviour could be generated have been presented, including the possibility that there is an independent 12.4 h circatidal oscillator, or that pairs of circalunidian (24.8 h) clocks run in antiphase (17,18). Evidence has been provided to support both viewpoints, sometime even on the same dataset. The problem with many of these studies is that there is no direct evidence at the molecular level to support one over the other competing theories. More recently, knockdown by dsRNAi of both *per* and *Clock* disrupted the circadian modulation of circatidal locomotor activity but not the circatidal period of the mangrove cricket, *Apteronemobius asahinai* (19,20). A similar result was observed in our previous study of the speckled sea louse, *Eurydice pulchra*, in which knockdown of Ep*per* to ∼20% of normal levels dramatically disrupted the circadian phenotypes of chromatophore dispersion, Ep*tim* mRNA cycling (the only canonical clock component that shows cycling in *Eurydice*), but left the circatidal locomotor cycle intact (12). Furthermore, maintaining the sea louse in constant bright light that severely damps the circadian cycles in chromatophore dispersion and Ep*tim* mRNA expression had no effect on the period or robustness of circatidal rhythms (12). Similar conclusions were reached with the circalunar spawning rhythms of the marine polychaete annelid *Platynereis dumerilii* in which pharmacological disruption of circadian clock components with CK1 inhibitors affected circadian molecular and behavioural rhythms but failed to impact the reproductive cycle (15). Consequently, it would appear that the circadian oscillator as a module does not contribute to core circatidal/circalunar function in at least three higher eukaryotes.

At odds with the conclusion reached above however, and unlike the case with circalunar cycles in *P. dumerilii*, treatment of *E. pulchra*, with the same CK1 inhibitors generated a dose-dependent lengthening of period for both free-running circadian and circatidal phenotypes (12). Given that CK1 modulates the stability of PERIOD (8-10) this result was intriguing given that direct dsRNAi knockdown of Ep*per* disrupted the circadian but not the circatidal mechanism (12). We suggested at the time that in *Eurydice*, either CK1 may have several targets (8,21) including an unknown ‘tidal’ protein, or that the inhibitors might be non-specifically disrupting the phosphorylation of an unknown tidally relevant kinase. *D. melanogaster* DBT also phosphorylates CLOCK (CLK) (22-24) and CLK stabilises CYCLE (CYC) (25). A further possibility is that the CK1 inhibitors might have disturbed EpCLK-BMAL1 mediated transcription (BMAL1 is homologous to CYC) but that EpCLK-BMAL1 are required independently for the expression of circadian or circatidal phenotypes.

In this study, we report that these CK1 inhibitors indeed affect *Eurydice* EpCLK-BMAL1 mediated transcription via the disrupted phosphorylation of both transcription factors, thereby implicating these positive regulators in the CK1-sensitive circatidal mechanism. We therefore directly targeted Ep*bmal1* with dsRNAi. In addition, we also knocked down the major *Eurydice* negative regulator Ep*cry2* (12). We observe that disruption of the former generates both circadian and circatidal phenotypes whereas Ep*cry2* knockdown predominantly affects only the circadian phenotypes. We therefore propose a neurogenetic model that explains how circadian and circatidal phenotypes are generated.

## RESULTS

### Casein kinase inhibitors inhibit Eurydice CLK-BMAL1 mediated transcription

CLK-BMAL1 heterodimers bind to E-boxes in *per* promoters to transcriptionally activate *per* (26,27) so we utilised the Drosophila S2 cell transcription assay where an E-box containing enhancer is fused to a luciferase reporter (12). Co-transfection of Ep*Clk/Epbmal1* gave high levels of luciferase activity (12) (Figure 1A) that were dose-dependently reduced (F_4,10_=249.7, p∼0) by adding the PF670462 CK1 inhibitor which is more selective for the CK1δ isoform in mammals (21)(Figure 1A). The inhibition of trans-activation occurred in the absence of EpPER (and endogenous *Dm*PER) suggesting that the inhibitors disrupt the phosphorylation of EpCLK-EpBMAL1. S2 cells were transfected singly with either tagged *EpBmal1* or *EpClk*, or co-transfected. Co-transfection revealed a number of additional higher molecular weight isoforms for each corresponding protein in Western blots (Figure 1B). Lambda alkaline phosphatase (λPP) restored each of the bands to their singly transfected original sizes. Administering PF6700462 revealed changes in *Ep*CLK mobility towards the hypophosphorylated isoforms (Figure 1B). Within each lane of the gel, the two *Ep*BMAL1 bands revealed an increase in the relative intensity of the higher MW isoform (Figure 1B red arrow) compared to the lower. This ranged from 47, 47 and 40% without the drug (lanes 3,4 and 9, red arrow, Figure 1B) to 59 and 56% when 5 μM of PF670 were added and 59% with 10μM (lanes 7, 11 and 8 respectively).

**Figure 1.**
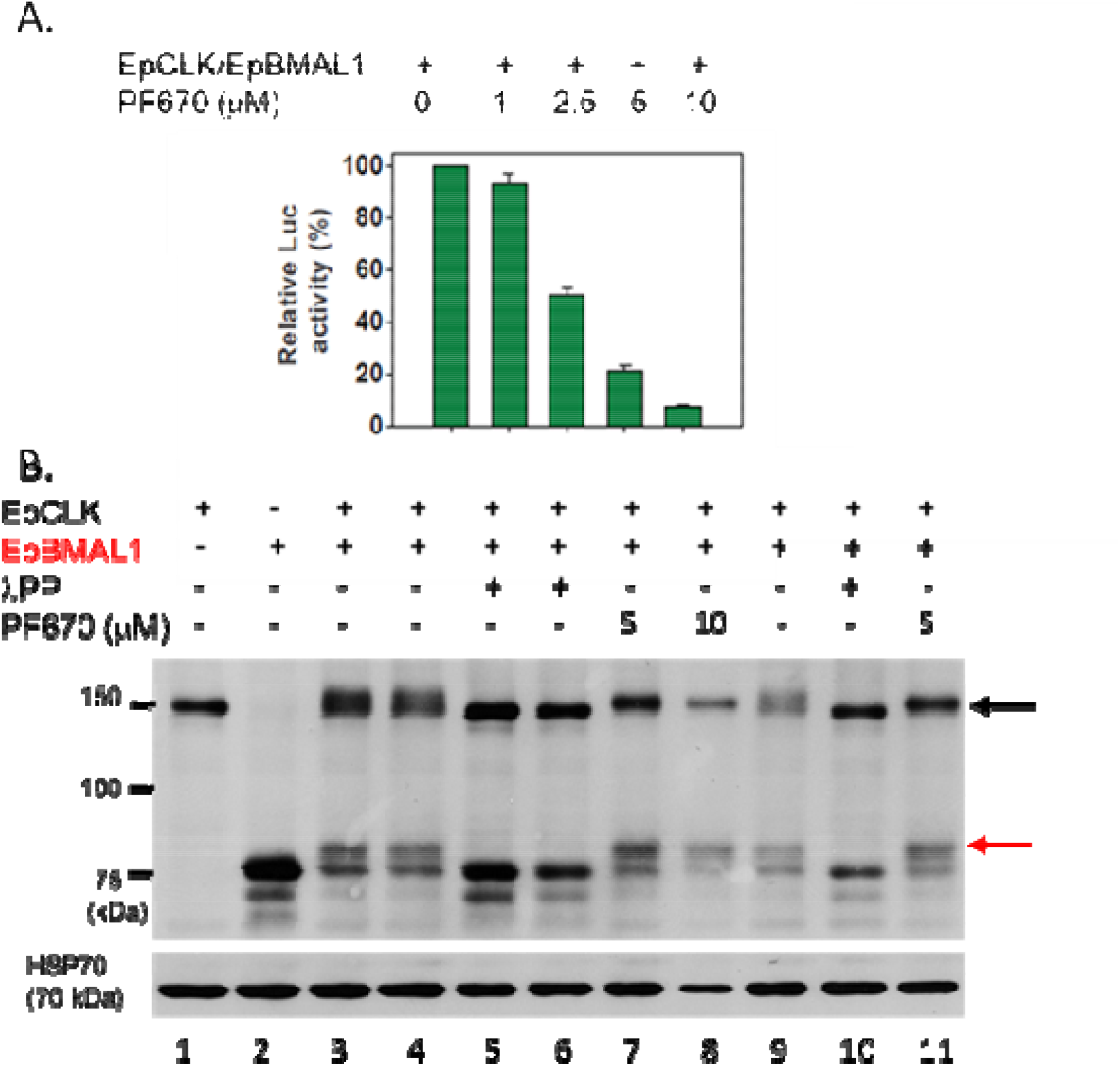
CK1ε/δ inhibitor PF670462 reduces EpCLK/EpBMAL1 E-box mediated transcription by modulating phosphorylation. A. PF670 represses E-box mediated EpCLK-BMAL1 mediated transcription in S2 cells (F_4,10_=249.7, p∼0, means + sem). B. PF670 alters the phosphorylation profiles of EpCLK (black arrow) and EpBMAL1 (red arrow) in Drosophila S2 cells, λPP lambda protein phosphatase (corresponding figures for PF480 in Figure S1).

We observed an almost identical dose-dependent reduction in transcriptional response in S2 cells with the CK1 inhibitor PF4800567 (F_4,10_=169.7 p∼0), which is more selective for CK1ε in mammals (21)(Figure S1A). In the corresponding western blot we obtained the same hypophosphorylation of EpCLK at doses of the inhibitor of 5 and 10μM (compare lane 3,4 without inhibitor to lanes 6 and 7). For EpBMAL1 we obtained a similar relative increase in intensity of the higher MW band (61 and 69% with inhibitor compared to 50 and 50% without, lanes 6, 7 compared to 3 and 4 (Figure S1B). Consequently, albeit in a heterologous system, these CK1 inhibitors at different concentrations show consistent effects on the phosphorylation profiles of both EpCLK and EpBMAL1, thereby implicating the positive regulators in the CK1 inhibitor-mediated lengthening of circatidal periods observed previously (12). We therefore tested for any effects of direct manipulation of EpCLK-EpBMAL1 on circatidal rhythmicity by using gene knockdown.

### dsRNAi knockdown of Ep*bmal1* and Ep*cry2* disrupts circadian phenotypes

We employed dsRNAi for *in vivo* knockdown of both the positive regulators and, in addition, the potent *Eurydice* negative regulator, Ep*cry2* (12). Exhaustive attempts to reliably reduce Ep*Clk* levels failed, but a consistent reduction of >50% for both Ep*bmal1* and Ep*cry2* transcripts was observed in preliminary experiments from the 3/4^th^ day after injection and maintained for several further days compared to controls injected with RNAi to yellow fluorescent protein *(WT*^*YFPi*^) (Figure S2A-E). Ep*cry2* and Ep*bmal1* mRNA levels in the control and knockdown animals over the circadian cycle revealed no circatidal or circadian cycling (12) but highly significant reductions to 44% and 43% of control *WT*^*YFPi*^ values were observed for the cognate transcripts respectively (Figure 2A, 2C). In gene-dosage terms, the dsRNAi generates animals that have less than the 50% that would be expected for individuals heterozygous for a wild-type and a null allele for both Ep*cry2* and Ep*bmal1*. Ep*tim* mRNA cycling is observed in Ep*bmal1* RNAi (Ep*bmal1i)* individuals compared to *WT*^*YFPi*^ during the 4^th^ day of DD but levels of Ep*tim* in Ep*bmal1i* were, as expected, significantly reduced to 71% of those in *WT*^*YFPi*^ (Figure 2B). The Ep*tim* mRNA cycle was dramatically damped in Ep*cry2i*, with overall transcript levels at 87% of those in *WT*^*YFiP*^ but there were no significant ANOVA effects due to the small number of replicates in this experiment (Figure 2D).

**Figure 2.**
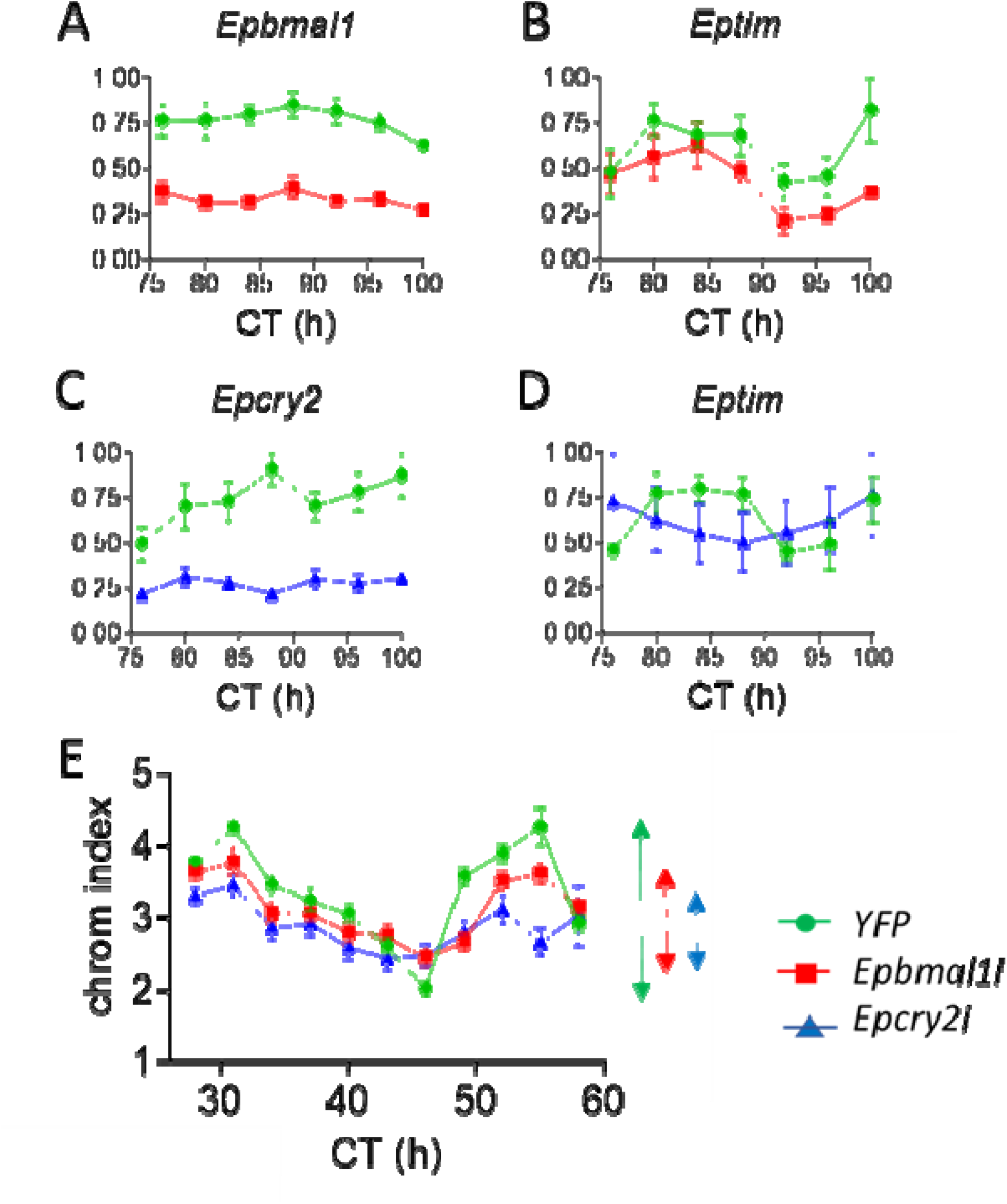
Knockdown of Ep*bmal1* disrupts circatidal and circadian phenotypes. A-E. dsRNAi of Ep*bmal1* (Ep*bmali*, red square) and Ep*cry2* (Ep*cry2i*, blue triangle) compared to WT^*YFPi*^ control (green circles). A, C. Ep*bmal1* and Ep*cry2* transcript levels are significantly reduced in *Epbmali* (n=6, F_1,58_ = 154.8), and Ep*cry2i* (n=4, F_1,34_ =10.38), Genotype p = ∼0 for both. There are no significant effects of Time. X-axis, circadian time (CT). Y-axis normalised relative abundance. Means +/- sem B, D. Ep*tim* cycles are present in *Epbmal1i* but levels of Ep*tim* are significantly reduced. (n=6, Genotype F_1,58_ = 11.1 p =1.5 × 10^−3^, Time F_6,58_ = 4.5 p=8 × 10^−4^). Ep*tim* cycles in Ep*cry2i* are altered but there are no significant effects by ANOVA (n=4). X-axis, CT; Y-axis normalised relative abundance. Means +/- sems E. Circadian chromatophore cycle. Peak-to-trough amplitudes shown with double-headed arrows on right of panel. ANOVA reveals significant effects for Time (F_10,772_=29.7) Genotype (F_2,772_ = 20.5) and G x T interaction (F_20, 772_=2.82, all p< 4 × 10^−5^). Dunnett’s *post hoc* tests reveals that both Ep*bmal1i* (p=0.0002) and Ep*cry2i* (p= 0.00004) are significantly different from WT^*YFPi*^ controls. X-axis, CT, Y-axis chromosome dispersion index

As injections *per se* can disrupt the circadian modulation of swimming episodes (12), we instead analysed the free-running circadian chromatophore cycle. Control animals (*WT*^*YFPi*^) exhibited a clear circadian cycle of chromatophore dispersion. We observed highly significant changes in the profiles of Ep*bmal1i and* Ep*cry2i* individuals compared to *WT*^*YFPi*^ controls that was reflected in the ANOVA and Dunnett *post hoc* tests comparing each experimental group with the control. The peak-to-trough amplitude index was reduced to ∼1-1.25 units for Ep*bmal1i* and Ep*cry2i* compared to 2.5 for *WT*^*YFPi*^ (Figure 2E). The experimental genotypes also showed a delayed upswing on the second DD cycle, particularly Ep*cry2i* (Tukey *post hoc* Ep*cry2i v* WT^*YFPi*^ p=0.00002, Ep*cry2i v* Ep*bmal1i* p=0.003, Figure 2E). Consequently, dsRNAi of both the positive and negative regulators was effective in altering circadian phenotypes.

### Ep*bmal1* but not Ep*cry2* knockdown reduces power and amplitude of circatidal behavioural cycles

To analyse the effects of gene knockdown on circatidal swimming, groups of animals were harvested in three main collections on spring tides during full and new moon in the 2016 season. They were injected with dsRNAi constructs after one day and then maintained under constant conditions for 4 days so that their free-running activity recordings were initiated at CT96 (see Methods). Locomotor activity histograms for each collection for the three knock-down ‘genotypes’ are shown in Figure 3. While the number of animals generating sufficient data for analysis is relatively small for each genotype within each collection (median = 21) overall the *Epbmal1i* animals reveal a less robust circatidal phenotype than the control WT^*YFPi*^ and the *Epcry2i* **(Figure 3A-C)**. More specifically, in the mid-summer collection (Figure 3A) clear circatidal 12.4 h peaks of locomotor activity are observed in both the WT^*YFPi*^ and Ep*cry2i* individuals. These activity peaks occur just after high tide when the animals emerge from their sand burrows to forage (vertical coloured lines in Figure 3 depict the alternate subjective night-time high tides). In Ep*bmal1i* individuals the pattern is far less clear, with a seeming mixture of circatidal and circalunidian (24.8 h) swimming cycles. A second summer collection (Figure 3B) reveals very clear high amplitude circatidal cycles for WT^*YFPi*^ as well as for Ep*cry2i* individuals. The Ep*bmal1*i animals also show circatidal cycles, but the activity troughs are less well-defined compared to the other two groups. Finally, for three late summer/early autumn collections when animals were less abundant, locomotor data was pooled (by synchronising activity to local tidal time). Locomotor activity levels were generally reduced and circatidal rhythms were less prominent compared to earlier in the season and no obvious inter-genotype effects were observed in these profiles (Fig 3C).

**Figure 3.**
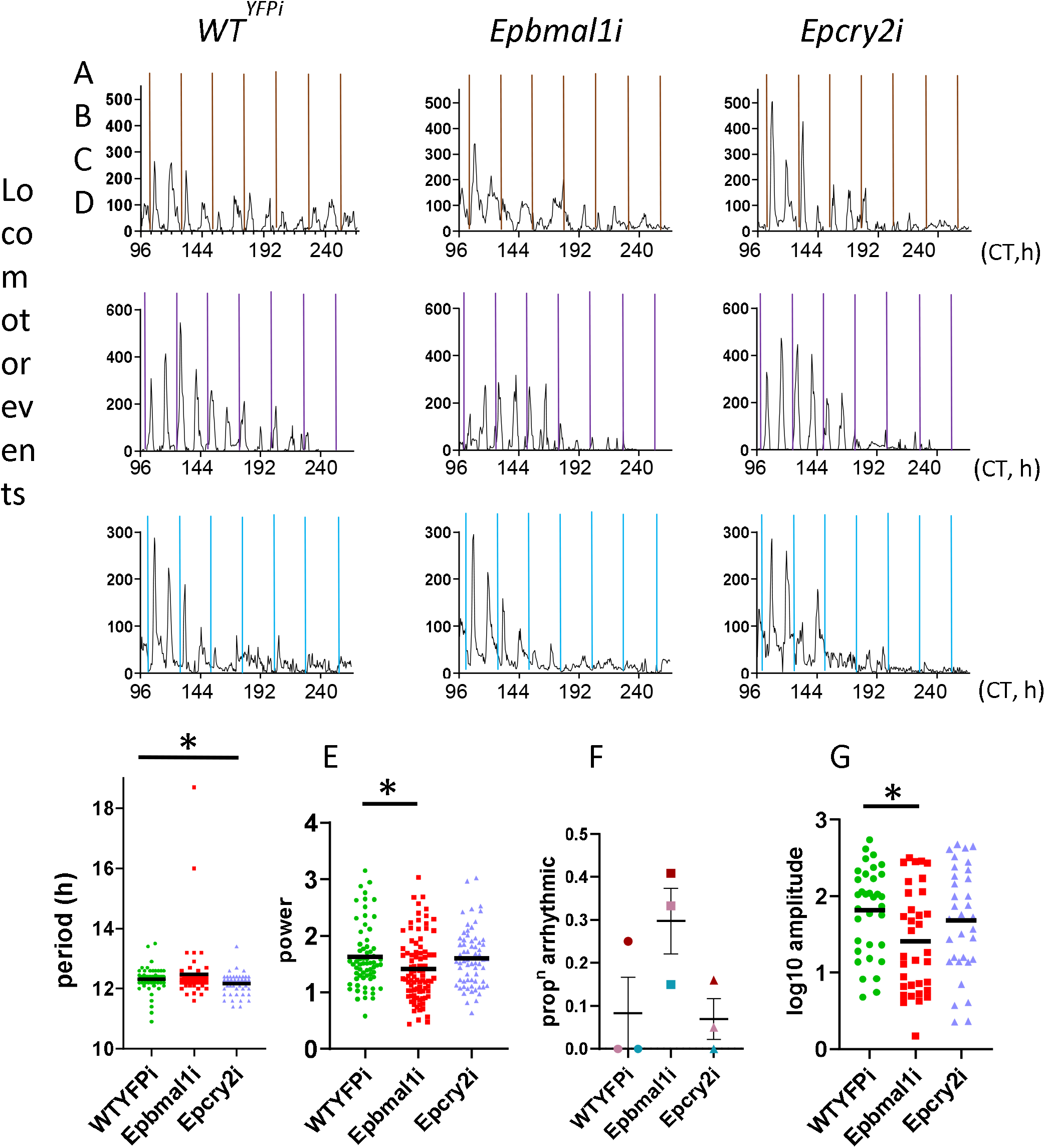
Circatidal rhythms in knockdown animals in season 1 (2016) A-C Histograms of mean activity in 30 min bins of animals collected during spring tides in mid-summer, full moon (A), midsummer new moon (B) and in late summer/autumn full moons (C). Standard errors of means are omitted for clarity. Panels on left are WT^*YFPi*^, middle are Ep*bam1i* and right are Ep*cry2i*. Vertical coloured lines represent the night-time high tide at Llanddona beach ie every 24.8 h. Animals were collected on the beach injected with dsRNAi constructs the following day and left for 3 days in DD before being placed in activity monitors to acclimatise for one further day in DD before activity recording started at CT96. For the new moon collection, B, a power cut truncated the time series. Ns for WT^*YFPi*^, Ep*bmal1i* and Ep*cry2i* are in A, 18, 21, 20, B, 23, 20, 20 and C, 24, 49, 25 respectively. Y-axis, locomotor (swimming) events per 30 min time bin. X-axis Circadian Time (CT, h) in constant conditions. D-G illustrates the statistical analyses of the results shown in A-C. D, circatidal period of the three genotypes. There is a marginal reduction in period of Ep*cry2i* (p=0.047, see text). E. Rhythm power shows a significant reduction for Ep*bmal1i* (p=0.03). Dotted black line represents power of 1, above which most animals show significant rhythmicity by spectral analysis (see Methods). F. The proportion of arrhythmic animals of each genotype in each collection. The colour of symbols correspond to the colour of the night-time high tide in each collection (A-C). Ep*bmal1* animals show a higher proportion of arrhythmic animals than WT^*YFPi*^ and Ep*cry2i* in every collection G. Amplitudes of the circatidal cycles from panels A-C. Ep*bmal1i* animals have significantly reduced amplitudes compared to WT^*YFPi*^ controls (p=0.02). * p<0.05. Means shown as horizontal black lines.

For the data collected for the full 2016 season, ANOVA revealed a significant effect on circatIdal period (F_2,176_=3.57 p=0.03, Figure 3D and Table 1). We removed the two obvious long period outliers in Ep*bmal1i* (Fig 3D) but still obtained a significant ‘genotype’ effect (F_2,174_=3.7 p=0.027). Dunnett *posthoc* test comparing each group to the WT^*YFPi*^ control revealed a marginally significant reduction in period for Ep*cry2i* (p=0.047). Analysis of the circatidal power with ANOVA also generated a significant ‘Genotypes’ effect (F_2,217_=3.64, p=0.028, Fig 3E and Table 1) with Dunnett’s test producing a significant reduction in power of Ep*bmal1i* compared to WT^*YFPi*^ (p=0.032) but not for Ep*cry2i* (p=0.95). This is reflected in Figure 3E where, in contrast to WT^YFPi^ and Ep*cry2i* animals there is a large cluster of Ep*bmal1i* animals that have a spectral power <1 and are classed as arrhythmic. In addition, there are a few animals that had spectral power slightly >1 but were not significant by autocorrelation (see Methods). Figure 3F shows that for each of the three collections (panels A, B and C indicated by brown, purple and turquoise vertical lines, and corresponding coloured symbols in Fig 3F), Ep*bmal1i* always showed a higher frequency of arrhythmic animals than the corresponding WT^*YFPi*^ and Ep*cry2i* animals. A Fisher exact test on the numbers rhythmic/arrhythmic for the season generated a highly significant χ2 value (21.7, df=2, p=<0.0001). 33% of Ep*bmal1i* animals are classed as arrhythmic (30/90) compared to ∼9% for WT^YFPi^ (6/65) and ∼8% for Ep*cry2i* (5/65, Table1).

**Table 1.**
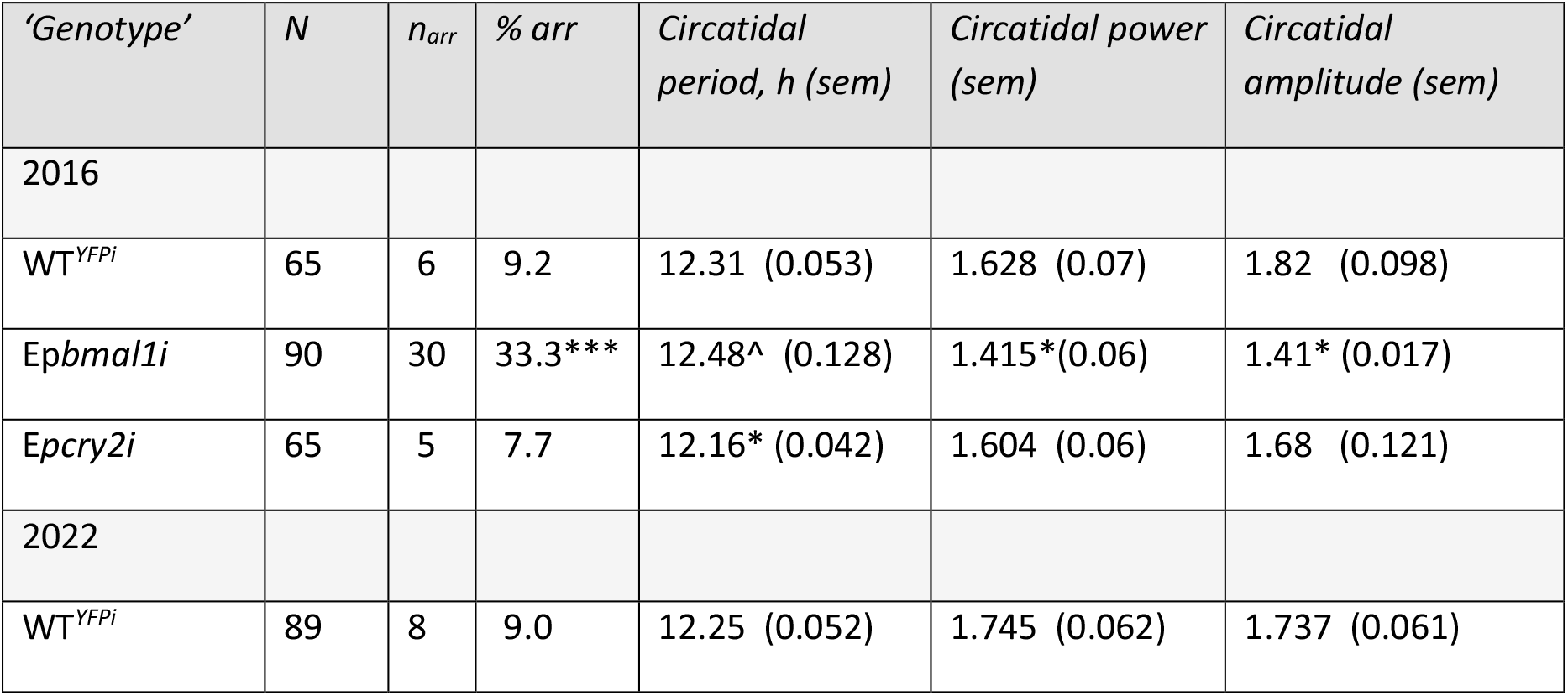
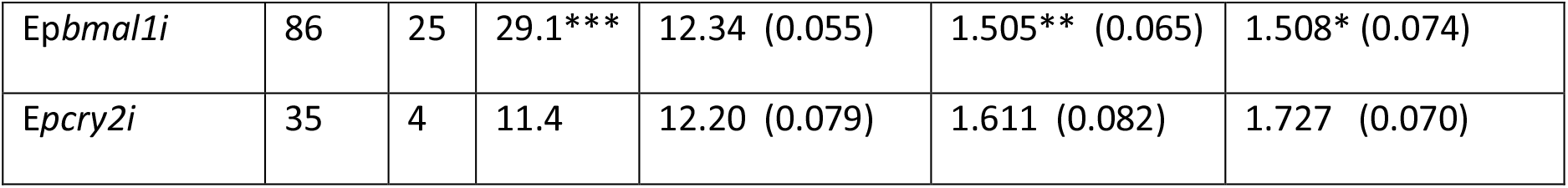
Circatidal results in two seasons of dsRNAi experiments. ***p<0.001, ** p=0.01, *p<0.05. The numbers arrhythmic were analysed by Fisher exact test (see text). ^The circatidal periods for the 2016 season include two very long period outliers for Ep*bmal1i*. If the two outliers are removed the period is 12.31 +/- 0.039 h (see text)

| ‘Genotype’ | N | $n_{arr}$ | % arr | Circatidal period, h (sem) | Circatidal power (sem) | Circatidal amplitude (sem) |
| --- | --- | --- | --- | --- | --- | --- |
| 2016 |  |  |  |  |  |  |
| $WT^{YFPi}$ | 65 | 6 | 9.2 | 12.31 (0.053) | 1.628 (0.07) | 1.82 (0.098) |
| *Epbmal1i* | 90 | 30 | 33.3*** | 12.48^ (0.128) | 1.415*(0.06) | 1.41* (0.017) |
| *Epcry2i* | 65 | 5 | 7.7 | 12.16* (0.042) | 1.604 (0.06) | 1.68 (0.121) |
| 2022 |  |  |  |  |  |  |
| $WT^{YFPi}$ | 89 | 8 | 9.0 | 12.25 (0.052) | 1.745 (0.062) | 1.737 (0.061) |
| *Epbmal1i* | 86 | 25 | 29.1*** | 12.34 (0.055) | 1.505** (0.065) | 1.508* (0.074) |
| *Epcry2i* | 35 | 4 | 11.4 | 12.20 (0.079) | 1.611 (0.082) | 1.727 (0.070) |

Figure 3G illustrates the amplitude of the cycle taken as a ratio of each peak value compared to the successive trough for each of the time series shown in Figs 3A-C. ANOVA of the log_10_ ratios generated a significant F-ratio (F_2,98_=3.46 p=0.03) and Dunnett’s test revealed a significant difference between WT^*YFPi*^ and *Epbmal1i* (p=0.02) but not between WT^*YFPi*^ and Ep*cry2i* animals (p=0.61). Consequently, Ep*bmal1i* circatidal rhythms are compromised by knockdown for amplitude, power and proportion of arrhythmic animals. Table 1 illustrates the results of the various parameters that were analysed.

### A second season of injections

These *in vivo* results suggest that *Eurydice* circatidal rhythms are more sensitive to reductions in Ep*bmal1* than to Ep*cry2*. We therefore repeated our behavioural experiments in the 2022 collecting season using the same dsRNAi constructs (Figure 4). Collections were made in early, mid and late summer with two harvested in the autumn (median number per genotype per collection =21). In the early summer, insufficient animals were collected so we focused on *WT*^*YFPi*^ and Ep*bmal1i* injections (Figure 4A). For the two autumn collections, numbers were also small so they were pooled to provide sufficient data for the same two knockdown genotypes (Figure 4D). The early summer collection (Fig 4A) reveals that the WT^*YFPi*^ animals are clearly circalunidian, with ∼25 h cycles of swimming activity with the peak a few hours after the nigh-time high tide. In contrast, the Ep*bmali* animals show a mixed profile of circatidal and circalunidian cycles. In the midsummer collection, WT^*YFPi*^ and Ep*cry2i* animals show very similar profiles, with circatidal cycles of activity that in the former coalesce into circalunidian cycles towards the end of the time series. The Ep*bmal1i* animals maintained a predominantly low amplitude circalunidian cycle (Figure 4B). The late summer animals reveal prominent circatidal cycles for both WT^*YFPi*^ and Ep*cry2i* groups throughout the time series. In contrast the profiles for Ep*bmal1i* are less well defined and fade badly in the second half of the experiment (Figure 4C). Finally, with the late autumn animals, which were maintained in LL after injection to observe whether different lighting regimes played any role in the phenotypic responses, WT^*YFPi*^ controls showed clear circatidal cycles whereas Ep*bmal1i* produced very low amplitude circatidal cycles for the first half of the time series that gradually became desynchronised from the night-time high tide (Fig 4D). Consequently, we observe very similar results with DD and LL as expected from our previous study, in which LL did not disturb the circatidal cycle (9).

**Figure 4.**
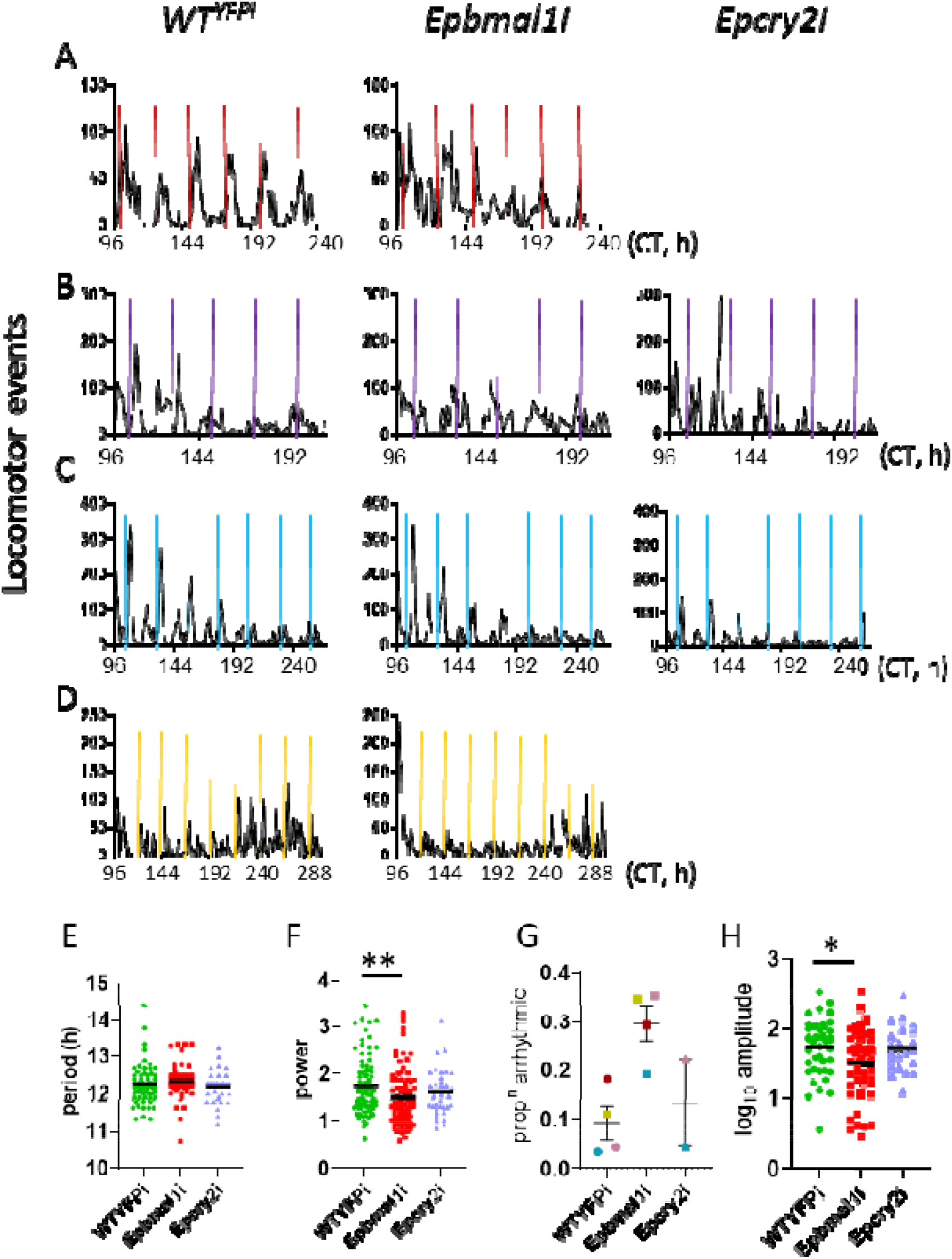
Circatidal rhythms in knockdown animals in season 2 (2022) As in Figure 3, but 4 collections from early summer (A), mid-summer (B), late summer (C) and autumn (D) where the animals were instead maintained in LL after injection. Ns for WT^*YFPi*^, Ep*bmal1i* and Ep*cry2i* are A; 22 and 17 (no Ep*cry2i* animals), B; 21, 17, 9 C; 28, 26, 21 and D; 18, 26, 5 respectively. No histogram is presented for the 5 Ep*cry2i* animals in D given the very low N. X-axis, circadian time (CT, h) in constant conditions. Y-axis, locomotor (swimming) events per 30 min time bin. E-H Statistical analyses of time series in A-D. E. There are no significant effects on period for the three genotypes. F. There is a significant reduction in power for Ep*bmal1* (p=0.01) with a large cluster of individuals arrhythmic with spectral power <1. G. Proportion of arrhythmic animals in each collection colour coded as in Figure 3. Ep*bmal1* animals show the highest frequency of arrhythmicity in each collection. H. Ep*bmal1* amplitude is significantly reduced compared to WT^*YFPi*^ controls (p=0.027). * p<0.05, **p=0.01. Means shown as horizontal black lines.

ANOVA of the full season data of circatidal periods did not generate a significant Genotype effect (F_2,170_=1.2, Figure 4E) whereas rhythm power (F_2,207_=3.77, p=0.025, Figure 4F) and rhythm amplitude (F_2,107_=3.64, p=0.03, Fig 4H, Table 1) were significantly affected by treatment. Dunnett tests revealed that *Epbmal1i* animals showed significantly lower rhythm power (p=0.01) and amplitude (p=0.027) than WT^*YFPi*^. This is reflected in the substantial number of Ep*bmal1i* animals showing a spectral power <1 compared to the other groups (Figure 4F) and the proportion of arrhythmic Ep*bmal1i* animals in each collection (Figure 4G). Fisher exact test on the seasonal numbers of arrhythmic versus rhythmic animals in each genotype was highly significant (χ2=13.3, df=2, p=0.001). 29% (25/86) of Ep*bmal1i* animals were arrhythmic, whereas in the WT^*YFPi*^ and Ep*cry2i* groups these values were 9% (8/89) and 11% (4/35) respectively (see Table 1).

Over both seasons of collections we have consistently observed that Ep*bmal1i* animals show significantly reduced circatidal rhythm power and amplitude that is also reflected in the significantly higher proportion of arrhythmic animals compared to both the WT^*YFPi*^ controls and the Ep*cry2i* group. While we also observed slightly longer circatidal periods in Ep*bmal1i* and slightly shorter periods in Ep*cry2i*, in the former, these differences were not significant, and in the latter, they were significant in the 2016 season, but not in 2022. Figure 5 illustrates the circatidal behaviour resulting from pooling the collections for each season and synchronising them to local tidal time. Ep*bmal1i* animals show relatively poorly defined circatidal rhythms compared to the other two groups in both seasons.

**Figure 5.**
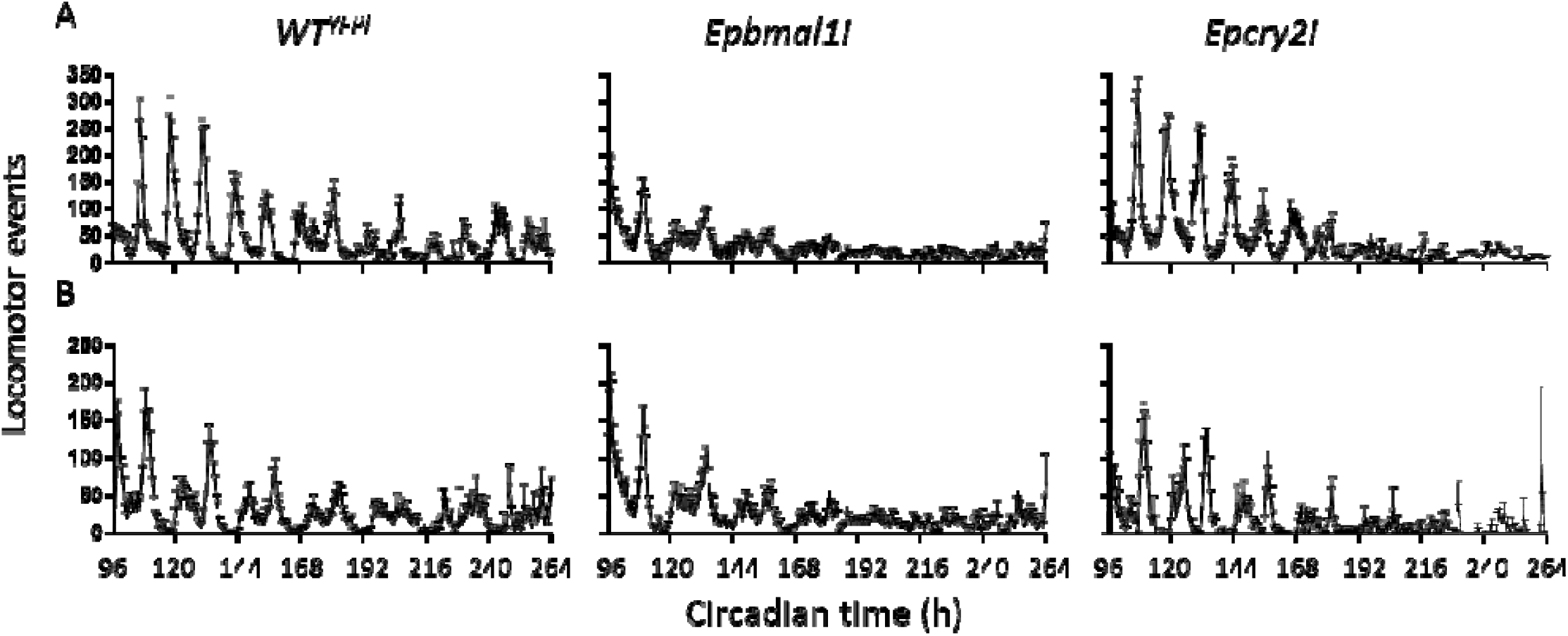
Circatidal rhythms of three dsRNAi genotypes. A. season 2016 B. season 2022. Collections were pooled for each season and synchronised to the local tidal time. X-axis, circadian time (h) in constant conditions. Y-axis, means +/-sem of swimming events per 30 min time bin.

## Discussion

The link between our previous observation of a dosage-dependent effect on circatidal periods by administering CK1 inhibitors (12) and our approach of knocking down EpBMAL1 and EpCLK, was forged by our results in the S2 cell transcriptional assay. We found that the CK1 inhibitors reduced transcription of an E-box-mediated reporter and that the phosphorylation profiles of EpCLK-EpBMAL1 were altered in the process. We cannot unambiguously state that the inhibitors were directly affecting the phosphorylation of the positive circadian regulators because it is conceivable that the inhibitors may be targeting another kinase non-specifically (21). Yet irrespective of the kinase identity, changes in the post-translational modifications of EpBmal1 and EpCLK by the inhibitors indirectly implicated these two positive circadian regulators in the expression of circatidal behaviour.

We therefore employed dsRNAi to knockdown the positive regulator Ep*bmal1*, and, as a counterpoint, also the negative regulator Ep*cry2* and were successful in reducing the gene dosage of both loci to <50%. While the knockdowns were not as effective as in our previous study with Ep*per*, where we obtained knockdown to ∼20% of wild-type levels (12), for both Ep*bmal1i* and Ep*cry2i*, the knockdowns were sufficient to generate significant changes in circadian phenotypes. From two seasons of collections, in 2016 and 2022, we obtained markedly similar behavioural results leading us to conclude that the circatidal swimming rhythm of *Eurydice* is more sensitive to reductions in Ep*bmal1* dosage than to similar reductions in Ep*cry2*. Indeed the Ep*cry2i* knockdown appeared to have a more dramatic effect on both the circadian cycles in chromatophore dispersion and Ep*tim* abundance, yet there was little evidence for any consistent effect on the Ep*cry2i* circatidal cycle. While there were marginal changes in circatidal period in both the Ep*bmal1i* and Ep*cry2i* these were neither significant for the former nor maintained statistically between seasons for the latter (Table 1). The most striking effect was on the robustness of the circatidal rhythm where in both seasons, Ep*bmal1* circatidal power and amplitude were significantly reduced, with 29-33% of this ‘genotype’ showing arrhythmicity compared to 6-11% of the WT^*YFPi*^ and Ep*cry2i* animals.

Both the Ep*bmal1i* and Ep*cry2i* animals reduced gene dosage to less than that of an animal heterozygous for a wild-type and null mutant allele. For comparison, heterozygous *bmal1/+* mice do not show a significant difference in free-running period nor amplitude compared to wild-type (28,29). In contrast Drosophila *cyc*^*0*^*/+* heterozygotes show a lengthening of free-running circadian locomotor period of 0.8 h compared to wild-type with no apparent differences in robustness as measured by the proportion of arrhythmic animals (30). Indeed, these Drosophila results with *cyc+/cyc*^*0*^ heterozygotes had encouraged us that our >50% knockdown might reveal a circatidal phenotype.

One caveat to our approach is that unlike gene knockouts, knockdowns do not reduce the targeted gene dosage to zero and so it could be argued that the circadian system in *Eurydice* is simply more sensitive to gene dosage disruptions than the circatidal phenotype. Perhaps then, further reduction in Ep*cry2* dosage below 43% might reveal a more striking circatidal phenotype. However, knockdown of Ep*per* to ∼20% of normal dose that obliterates both the Ep*tim* mRNA and the chromatophore circadian cycles, did not alter circatidal behaviour (12). Nevertheless, it is possible that a complete knockout of Ep*cry2*, Ep*per* or Ep*tim* might generate a circatidal effect. Unfortunately, the long and complex life cycle of *E. pulchra* (31) in addition to our inability to rear it in the laboratory makes a CRISPR/Cas9 gene editing approach impractical. The main advantage to using dsRNAi in the adult is that possible confounding developmental defects of a gene knockout are avoided. In *Drosophila, cyc*^*0*^ (and *Clk*^*jrk*^) mutants show reductions in the numbers of pacemaker LNv clock neurons in both adults and larvae as well as abnormal projections from these cells (32,33).

Our current knockdown results with Ep*cry2* and those of our previous study with Ep*per* (12) (that also disrupts the Ep*tim* mRNA cycle) generate circadian but not circatidal phenotypes, whereas manipulation of Ep*bmal1*, affects both types of rhythm. It is therefore tempting to speculate that the negative circadian regulators (Ep*per*, Ep*tim*, Ep*cry2*) may not be involved in the generation of circatidal oscillations, whereas the positive regulator, Ep*bmal1* (and possibly Ep*Clk*) play more fundamental roles. A simple model would have dedicated but separate circadian and circatidal neurons in which all the canonical circadian clock molecules are expressed in the former cells, with cycling Ep*tim* expression driving the negative feedback loop. In the latter cells, the positive regulators EpBMAL1-EpCLK would be present, but not the negative regulators whose place would be taken by a novel circatidal regulator, whose expression would cycle with a ∼12 h period and would engage the circatidal negative feedback loop. The two oscillators would interact at the level of output given that *Eurydice* circatidal behaviour can show circadian modulation (9). The model can incorporate a circatidal cycling kinase/phosphatase element driven by the circatidal clock that would feed back onto EpBMAL1-EpCLK function and support cycling transcription of the novel negative circatidal regulator. This would only occur in circatidal cells, thereby maintaining the separate integrity of the two oscillators.

This model can be tested by extensive investigation of the expression patterns of canonical clock transcripts and proteins both temporally and spatially in the *Eurydice* brain. EpPER appears to be located in a pair of dorsolateral neurons that show circadian cycles of abundance and a further lateral cell that does not (12). These EpPER neurons are separate from those in *Eurydice* that express PDH (34) which is a marker for brain pacemaker cells in Drosophila (35). We are currently studying the expression of other clock components with a number of homospecific anti-sera that we have targeted against positive and negative regulators. Our model predicts that groups of neurons expressing EpPER, EpCRY2 and EpTIM and EpCLK-EpBMAL1 would represent circadian clock neurons, and neurons expressing the latter positive regulators, but not the negative regulators would define the circatidal neurons.

In conclusion, our results reveal Ep*bmal1* to be the first gene to be implicated in driving circatidal behaviour. Until now, the literature had reported only the genes that were not required to generate lunar-mediated cycles. From an evolutionary perspective, re-using components for both circatidal and circadian mechanisms would appear to be a pragmatic solution to solving two similar timing problems.

### Note added

While we were preparing the manuscript we became aware of similar work to ours by Kwiatkowski et al (36). This study develops the amphipod crustacean *Parhyale hawaiensis* as a circatidal model and uses gene editing to knock out the Ph*bmal1* gene. The results reveal that circatidal cycles of behaviour are obliterated with this null mutant. These results are consistent with ours. We thank Patrick Emery for allowing us to see the ms.

## Materials and Methods

### Animal collections and behavioral and chromatophore analyses

Animals were netted from Llanddona Beach, Anglesey, North Wales, UK at high water on spring tides (https://www.tidetimes.co.uk/menai-bridge-tide-times-20220620), from June to October and maintained in seawater in LD12:12 at 16°C. Swimming activity was recorded in constant darkness (DD) using LAM10 locomotor activity monitor (Trikinetics Waltham MA) at 16°C. The late autumn collection of 2022 was maintained in LL but the results were similar to those in DD (see Figure 4D). Animals that showed at least 5-6 tidal cycles and at least 600 beam interruptions over 7 days were included for further evaluation, except for the truncated collection in 2016 (see Figure 2B) where the criterion was reduced to 500 activity counts. Animals that died during the experiment generate a series of ‘0’ counts for the rest of the recording and these were removed. A few animals showed no activity at the beginning of the experiment but then started moving so these ‘0’ counts at the beginning of the record were also removed. Each time series (in 30 min bins) was analysed independently with spectral analysis and autocorrelation (37) (except for the truncated collection shown in Figure 3C where we had only 5 days of data due to a power cut-here we used a minimum of 500 activity counts as a threshold for further analysis).

An animal was considered rhythmic if the two independent time series were significant and generated a consistent period. For the spectral analysis significance meant that the peak in the spectral plot using the CLEAN algorithm was above the 99% confidence limits that were determined using 100 random iterations of the original data. In addition, a significant autocorrelation had also to support the period observed in the spectral plot (statistical protocols described in ref 33). The power of the rhythm was estimated by taking the peak value of the spectral density plot and dividing by the value of 99% confidence limit. Consequently a rhythmic animal has a power of >1, but only if confirmed by a significant autocorrelation. In most animals, the major peak in the spectral plot was in the circatidal range between 12 -13 h but the doublet at 24-26 h was also commonly significant. When the power of the doublet was greater than that of the singlet the circatidal period was taken as half of the doublet whereas the power taken was that of the doublet. In the 2016 season collections, 15/220 (6.8%) animals whose activity met the criteria for analysis had doublet>singlet periods whereas in 2022 the doublet>singlets group comprised 69/212 (32.5%). In one particular collection in early summer 2022 (see Fig 4A), 23/39 (59%) animals had doublet>singlet. Taking half the doublet as the circatidal period rather than the singlet period has no effect on the overall circatidal period (because the doublet is usually exactly twice the singlet) but it has an effect on the power calculation. In 2022 38%, 31% and 20% of the WT^*YFPi*^, Ep*bmal1i* and Ep*cry2i* animals had doublet>singlet periods, so they are distributed roughly equally among genotypes (Fisher exact χ2 = 2.6, df=2, p=0.27).

The amplitude of the circatidal cycle was taken by generating an average activity plot based on mean activity events every 30 min for each ‘genotype’ for each collection. The activity value at each peak of the cycle was divided by the value at the next trough. If the 30 min bin corresponding to the trough (the one with the lowest mean activity level) showed a mean activity level <1 locomotor event, a value of 1 was used to avoid generating enormous ratios. The amplitude for each peak/trough was taken for each time series for each genotype for each collection. Circatidal/circalunidian peaks were time-matched among the three genotypes within each collection. This generated between 9-16 amplitude values per time series. Transformation of peak/trough ratios to log_10_ was required to equalise variances among the genotypes before ANOVA. All animals were analysed ‘blind’ to their knockdown genotype.

To assess chromatophore rhythms, animals were snap frozen in liquid nitrogen at defined tidal and circadian times, chromatophore patterns imaged by digital camera and scored ‘blind’ using the Hogben and Slome 5-point index (38) which was modified to include 0.5 point scoring intervals (12). These animals were maintained in LD12:12 for 3 days post dsRNAi injection after which they were placed in DD, and scored every 3 h DD for 30 h during day 4-5 (from the second day of DD). Between 4-6 animals were scored for each time point from each collection. Heads were cropped and snap frozen for later qRT-PCR.

### dsRNAi

Double-stranded RNA (dsRNA) molecules of Ep*Clk*, Ep*bmal1* and Ep*cry2* were designed with the E-RNAi web-service (39) and synthesised by using a MEGAscript RNAi kit (Ambion, UK) (Table S1). For the dsRNAi control, the yellow fluorescent protein (YFP) gene from pEYFP-N1 (Clonetech, UK) was used. 200-250ng of dsRNAs was injected into the hemocoel using air pressure microinjection via glass microcapillary (40). Gene suppression was assessed by real-time quantitative RT-PCR. Ep*bmal1i* animals tolerated the injections rather better than the other two genotypes, so numbers surviving to full analysis were greater (see text).

The full length of Ep*Clk* (NCBI: KC885973), Ep*bmal1* (NCBI: KC885968) and Ep*cry2* (NCBI: KC885970) subcloned in pAc5.1/V5-hisA vector (12) were used as templates for the target sequences amplification (600bp for *Clk*, 587bp and 570bp for *bmal1*-1 and *bmal1*-2, respectively, 650bp for *cry2*). PCRs were primed using oligonucleotides containing a T7 phage promoter region (Table S1, below). Single stranded cRNA in both directions was synthesised and complementary RNA strands were hybridised and purified according to the manufacturer’s instructions. Double stranded products were analysed in agarose gels and concentrated by ethanol precipitation to 3µg/µl in nuclease-free water, aliquoted and kept at -80°C until use. For the dsRNAi control, the yellow fluorescent protein (YFP) encoding gene from pEYFP-N1 (Clonetech, UK) was used to generate a 400bp dsRNA with the same method described as the target sequences. Double-stranded RNA mixed with equal volume of 2x injection buffer (0.2mM sodium phosphate buffer pH 6.8, 10mM KCl) containing filtered food colour was injected into the haemocoel between anterior tergites through a glass microcapillary and using compressed nitrogen/air delivered by a PV830 PicoPump (World Precision Instruments, Inc). Animals were immobilised by leaving on ice and then transferred onto the ice-cold aluminium block using a sieve/mesh for injection under a microscope. About 130-160nl (200-250ng) of dsRNA was injected in each animal. Injected animals were placed on tissue for 2-3 minutes to ensure injected fluids did not leak out of the puncture wound (40). Gene suppression was assessed by qRT-PCR.

Initial collections for attempting dsRNAi and follow-up qPCR for Ep*Clk*, Ep*cry2*, and Ep*bmal1* were made in 2014 and 2015. Once the conditions were optimised, the dsRNAi experiments with Ep*bmal1* and Ep*cry2*, were performed from collections made from June to September 2016 and repeated from July to October 2022 (in 2020 and 2021 the pandemic had prevented us from travelling to Wales to harvest the animals). Animals collected from the beach were transfered immediately to Leicester where they were injected the following day with dsRNAi constructs (*WT*^*YFPi*^, Ep*bmal1i, or* Ep*cry2i*). The injected animals for locomotor recordings were then placed in DD for 3 further days, then placed in activity monitors for another day in DD to acclimatize, before the Trikinetics monitors were switched on (again in DD). Consequently, the activity recordings begin on the 5^th^ day after injection. A relatively small number of animals survived the immediate injections and of those about 40% did not provide behavioural data for more than 5 circatidal cycles and 600 activity events. Nevertheless, we were able to produce behavioural data for 65-95 animals per injected genotype in each season (except for Ep*cry2i* in season 2022, n=35) that met the selection criteria for time series analyses.

### Real-time quantitative RT-PCR

We examined the expression of Ep*bmal1*, Ep*Clk* and Ep*cry2* by qPCR to study the extent of the knockdown as well as to observe the effects on the Ep*tim* mRNA circadian cycle (12). Animals were maintained for different numbers of days in DD for the dsRNAi to take effect and initially mRNA was harvested from 5 animals, 10 h into subjective day (CT10, CT34, CT58 etc) at each day (Figure S2A). The Ep*tim* mRNA circadian cycle of expression was quantified by qPCR during the 4^th^ day of DD by taking samples every 4 h.

Total RNA was extracted from pooled heads using Trizol Reagent in conjunction with the PureLink RNA Mini Kit (Invitrogen). DNA contamination was removed by on-column PureLink DNase treatment (Invitrogen). 0.5-1µg total RNA from each sample was used for cDNA synthesis by using the High Capacity cDNA Reverse Transcription Kit (Applied Biosystems) and oligo_dT (12-18) primer (Invitrogen) in a 20µl total reaction volume for 120 minutes at 42°C and the reaction was terminated by heating to 85°C for 5 minutes.

Quantitative PCR was performed on the Roche LightCycler 96 instrument by using GoTaq qPCR Master Mix (Promega) with 1µl cDNA template and 0.5µl of 10mM each primer in a total volume of 25µl reaction. The cycling conditions were as follows: 95°C for 120 seconds, 40 cycles of 95°C for 15 seconds and 60°C for 60 seconds, and then followed by melting curve reaction at 95°C for 10 seconds, 65°C for 60 seconds and 97°C for 1 second. The primer pair for each gene was designed to amplify 100-130bp PCR products (Table S1). The relative quantification method from Roche LightCycler software was used to calculate gene expression and ratio error. The Standard curves were obtained using decimal dilution series of plasmid DNA. Transcript levels were normalised to the *Eurydice* ribosomal protein L32 gene *(RPL32)* and for Ep*tim* mRNA experiments, values were scaled to the timepoint with the highest expression level in *WT*^*YFP*^ controls to allow pooling of biological replicates and statistical analysis by ANOVA. *bmal1*-1 and *bmal1*-2 sequences were equally efficient for knockdown (see Suppl Fig 2C-E) and so *bmal1-1* was used for the injections.

### CK1 inhibition in S2 cells

The *Drosophila* S2 cells (Invitrogen) were maintained in HyClone SFX-insect medium (Thermo Scientific) supplemented with 10% fetal bovine serum (FBS) and penicillin-streptomycin antibiotics at 25°C as described previously (12). Cells were transfected with expression constructs by using Cellfectin (Invitrogen) according to the manufacturer’s instructions. Ep*Clk*, Ep*bmal1*, were amplified from their corresponding plasmids and sub-cloned into the *Drosophila* S2 cell expression vector pAc5.1/V5-HisA (Invitrogen) as reported previously (12). Control transfections, including only reporter construct and empty vector (pAc5.1/V5-hisA) established baseline activity. Luciferase activity was measured using the Dual Luciferase Reporter Assay Kit (Promega) and was normalised for transfection efficiency using a *Renilla* expression plasmid. At least three independent transformations were performed for each assay.

CK1ε/δ inhibitor, PF670 or PF480 (Tocris Biosciences) solution was added into S2 cells after 5-6 h transfection to a final concentration as indicated and the drug treated cells were incubated for 48 h at 25°C before harvest for luciferase activity (12) or western analysis. The lambda protein phosphatase treatment is described in the western blot analysis below.

### Western blot

Transfected cells were washed with ice-cold PBS, pelleted at 4°C and lysed in the RIPA buffer (Sigma) along with complete protease inhibitor cocktail (Roche) and PhosSTOP Phosphatase Inhibitor (Roche). For the protein phosphatase treatment, cells were lysed as described above, except in the absence of phosphatase Inhibitor and incubated with 400u lambda protein phosphatase (New England Biolabs) at 30°C for 1 h. About 50ug total protein from cell extracts were blotted and hybridised with the primary antibody of Mouse anti-V5 (Invitrogen) for CLK and BMAL1 expression then the secondary antibody of horseradish peroxidase-conjugated either anti-mouse or anti-rabbit IgG antibody (Sigma). Chemiluminescence detection was performed by using ECL Western Blotting detection Reagent (GE Healthcare). HSP70 was used as a general loading control but was not used in the quantification of the BMAL1 hypo/hyperphosphorylated isoforms which were quantified relative to each other using ImageJ software. Three different gels were run for the S2 cell westerns (biological replicates) with multiple lanes carrying both technical but also biological replicates, the latter treated with different drug doses.

## Data and materials availability

All raw data will be deposited in Dryad repository on acceptance (https://datadryad.org/stash).

## Acknowledgements

CPK SGW and DCW acknowledge BBSRC grants BB/E000835/1, BB/K009702/1 and BB/R01776X/1. This work was also supported by the Medical Research Council, as part of United Kingdom Research and Innovation (also known as UK Research and Innovation) [ *MRC File Reference No. MC_U105170643* ] to MHH. For the purpose of open access, the MRC Laboratory of Molecular Biology has applied a CC BY public copyright licence to any Author Accepted Manuscript version arising.

## Author contributions

The study was conceived and supervised by CPK, SGW and MHH. Funding was obtained by CPK, SGW and DCW. LZ and DCW collected the animals, LZ performed all the molecular and behavioural work. EWG wrote the scripts for statistical analyses of the data. LZ and DCW performed the chromatophore experiment and CPK performed the statistical analyses of chromatophore, qPCR and behavioural results. CPK wrote the draft ms and all authors contributed to the final version.

## Competing interest declaration

Authors declare no competing interests

## Notes

### Competing Interest Statement

The authors have declared no competing interest.

