## Supplemental Figures 1 and 2 and Table S1 for "The circadian clock gene *bmal1* is necessary for co-ordinated circatidal rhythms in the marine isopod *Eurydice pulchra* (Leach)"

**Figure S1. CK1ε/δ inhibitor PF480 reduces EpCLK/BMAL1 E-box mediated transcription by modulating phosphorylation**

A. PF480 represses E-box mediated EpCLK-BMAL1 mediated transcription in S2 cells ($F_{4,10}$=169.7 p~0, means + sem).

B. PF480 alters the phosphorylation profiles of EpCLK (black arrow) and EpBMAL1 (red arrow) in Drosophila S2 cells, λPP lambda protein phosphatase. Relative intensities of the two EpBMAL1 isoforms were quantified with Image-J software within each lane, so did not require running a HSP-70 loading control.

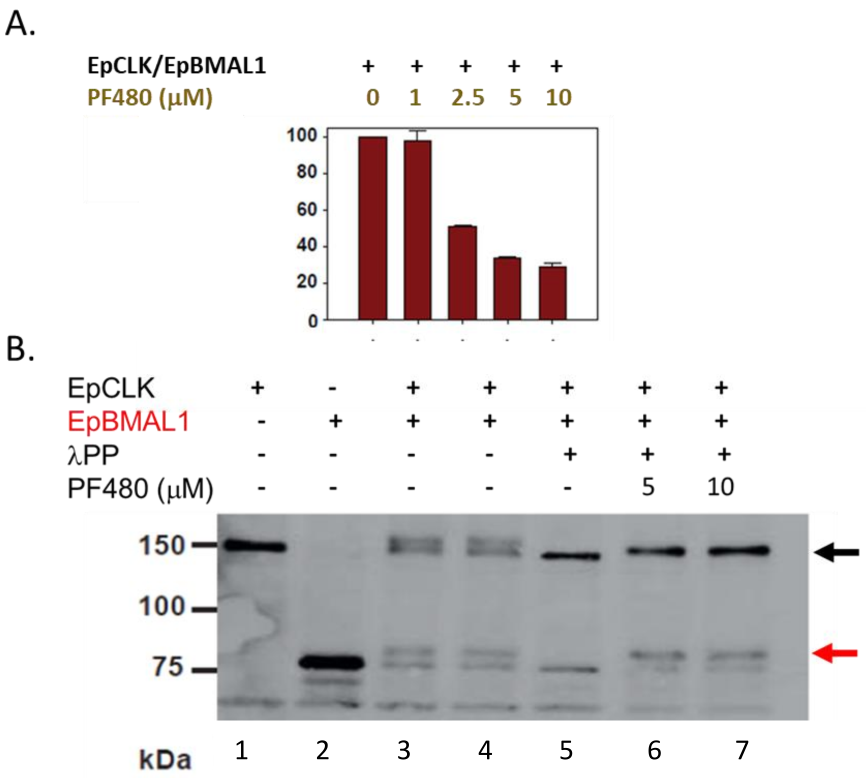

**Figure S2. Preliminary dsRNAi knockdown experiments for Ep*Clk*, Ep*bmal1* and Ep*cry2***

A, B. Two experiments attempting to knockdown Ep*Clk*. qPCR performed up to 6 days after injection. No consistent long-lasting knockdown was achieved. Means +/- ratio error was calculated using Roche LightCycler software

C, D. Two experiments using two different sequences of Ep*bmal1* (F1 and F2) show significant knockdown from days 4-9.

E. Knockdown of Ep*bmal1* (F1 sequence) and Ep*cry2* shows significant knockdown at days 3 and 4 after injection (see also Figure 2A-D). Data from combing separate experiments on *Epbmal1F1* and Ep*cry2* knockdown by setting YFP values to 100%.

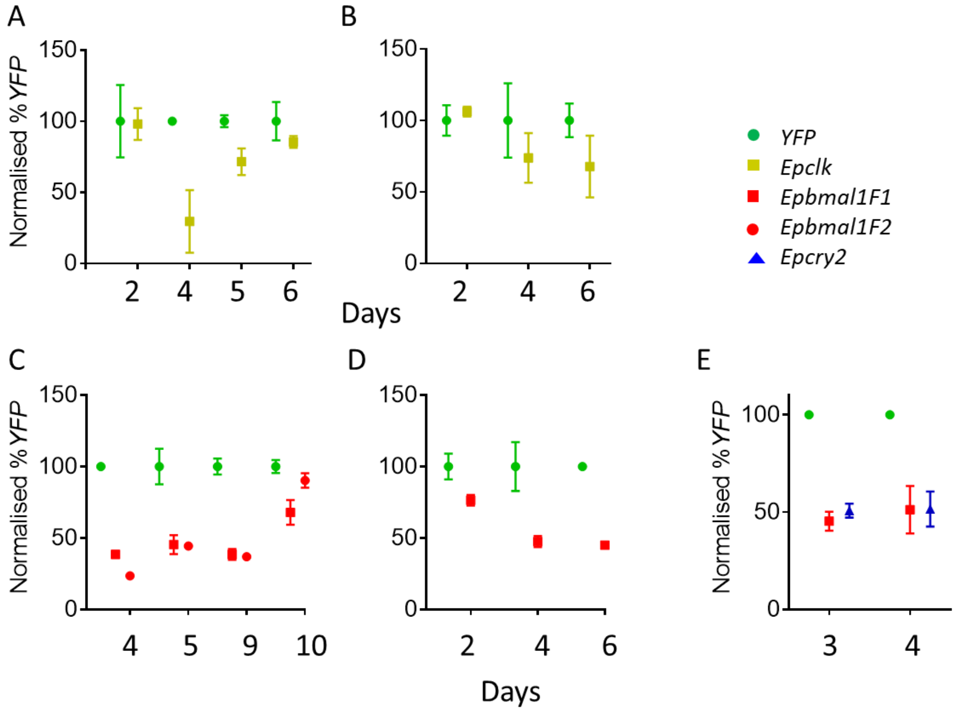

**Table S1 Primers for dsRNAi and qRT-PCR**

| | **Primer name** | **Forward sequence (5’→3’)** | **Reverse sequence (5’→3’)** |
|---|---|---|---|
| dsRNAi | *EpClk dsRNA* | *T7-CTTACTCCGTACCGCTCGTC* | *T7-GCTGCTGTTGCATGTGACTT* |
| | *Epbmal1_1 dsRNA* | *T7-GGTATCCGATTGTGCGTCTT* | *T7-AGAAGTCGACAGGTTCCCCT* |
| | *Epbmal1_2 dsRNA* | *T7-CCGTTCCAAAAGACACCACT* | *T7-ACTCTGGTCTCCTGTGTGGG* |
| | *Epcry2 dsRNA* | *T7-TCCAGATTCCGTGGGATAAGA* | *T7-CTCTCTTGTCCTTCTTCGGG* |
| | *yfp dsRNA* | *T7-AGGACGACGGCAACTACAAG* | *T7-GTCCATGCCGAGAGTGATCC* |
| qRT-PCR | *EpClk qPCR (122bp)* | *GCAACAGCAGACCTTCCTTC* | *GGGTGAGAGGGTTGAGAGGT* |
| | *Epbmal1 qPCR (131bp)* | *CTCTTCGTCGTTGGTTGTGA* | *GCCAGATCTTTCGGATGAAG* |
| | *Epcry2 qPCR (100bp)* | *ACCTGCCCAATCCTGTGTGG* | *GGCCTCCCAAACGAAGCTACC* |
| | *EpRPL32 qPCR (107bp)* | *CAAAATTGGAGGAAGCCAAA* | *TGCTTTGTTTTCTTGGCTGA* |
| | *Eptim3 qPCR (120bp)* | *AGCTGAATTTCCACCTCTGCG* | *CCAGAGTCGCGTTCCTCTTC* |

*T7- TAATACGACTCACTATAGGGAG(A)*
